# Mechanisms for plastic landmark anchoring in zebrafish compass neurons

**DOI:** 10.1101/2024.12.13.628331

**Authors:** Ryosuke Tanaka, Ruben Portugues

## Abstract

Vision is a sensory modality particularly important for navigation, as it can inform animals of their current heading (i.e. visual landmarks) as well as its changes (i.e. optic flow). It has been shown that head direction (HD) neurons in various species incorporate the visual cues into their heading estimates. However, circuit mechanisms underlying this process remain still elusive in vertebrates. Here, we asked if and how the recently identified HD cells in the larval zebrafish – one of the smallest vertebrate models – incorporate visual information. By combining two-photon microscopy with a panoramic virtual reality setup, we demonstrate that the zebrafish HD cells can reliably track the orientation of multiple visual scenes, exploiting both landmark and optic flow cues. The mapping between landmark cues and the heading estimates is idiosyncratic across fish, and experience-dependent. Furthermore, we show that the landmark tracking requires the lateralized projection from the habenula to the interpeduncular nucleus (IPN), where the HD neuron processes innervate. The physiological and morphological parallels suggest that a Hebbian mechanism similar to the fly ring neuron is at work in the habenula axons. Overall, the observations that the hindbrain HD cells of the larval zebrafish lacking an elaborate visual telencephalon shed new light on the evolution of the navigation circuitry in vertebrates.

## Introduction

Neurons whose activities reflect spatial relationships between animals and their environments have been identified in the brains of diverse animal species [1], likely supporting navigation. A simple example of those space-tuned cells is head direction (HD) cells, which fire when animals face a particular direction [2]. Because the direction of animals’ heading cannot be typically directly conveyed by sensory inputs, HD cells need to integrate the history of rotational movements animals make, a process called angular path integration. As a simple, yet biophysically plausible mechanism to implement the angular path integration, a dynamical model called the ring attractor has been proposed [3–5]. These ring attractors typically consists of neurons arranged on a topological ring, which excite nearby neurons while inhibiting far away ones. This connectivity gives rise to a single, stable bump of activity on the ring, which can be used to represent the head direction.

More than two decades of research have identified HD cells in various mammalian brain regions[6]. Yet, it has remained inconclusive in which region the tuning to head directions first emerges through ring attractor-like dynamics. Recently, a study on the larval zebrafish identified a group of GABAergic HD cells in the anterior hind-brain (aHB) [7]. This GABAergic aHB nucleus is likely homologous to the mammalian dorsal tegmental nucleus [8], one of the basal-most brain regions with HD cells in rodents [9, 10]. The dendrites and axons of these zebrafish HD cells form topographically organized columns in the dorsal interpeduncular nucleus (dIPN), such that cells tuned to the opposite head directions would inhibit each other – a key connectivity motif of the ring attractor. These observations offered a detailed circuit-mechanistic explanation of how head direction representation is generated in the vertebrate brain. In addition to the history of turning commands, animals can also utilize visual cues, such as landmarks and optic flow, to improve their sense of heading [11, 12]. In rodents, cortical visual areas (e.g., the retrosplenial cortex) are thought to provide visual information to HD neurons [13]. However, unlike mammals, the larval zebrafish lacks an elaborate visual telencephalon. This raises the possibility that there exist evolutionarily older, non-telencephalic pathways that route visual information to the HD neurons.

In the present study, we asked if and how the larval zebrafish HD neurons utilize visual cues. Using a two-photon microscope with an immersive virtual reality (VR) setup, we first demonstrate that the larval zebrafish HD cells can keep track of the animal’s heading relative to visual scenes. By manipulating the VR, we show that the HD neurons track the scene orientation exploiting both visual landmarks and optic flow cues. In addition, we find that the mapping from visual scenes to head direction neurons is idiosyncratic, and can be remapped in an experience-dependent manner. Finally, we demonstrate that inputs from the visual side of the habenula are required for anchoring the head direction representation to visual landmark. Our results demonstrate how a conserved circuit in the larval zebrafish brain can integrate multimodal information to robustly encode the heading direction, shedding new light to the evolution of spatial cognition in the vertebrates.

## Results

### GABAergic aHB cells keep track of the visual scene orientation

Recently, we identified a group of gad1b+ HD neurons in the aHB rhombomere 1 (rh1) of the larval zebrafish [7]. The preferred headings of these HD cells are topographically arranged, such that when the fish turns rightward, the bump of neuronal activity moves counterclockwise, as viewed from the top (Figure 1a). However, unexpectedly, the study did not detect any effect of visual feed-back on the bump movements [7]. We reasoned that this might be due to the fact that the visual stimuli were presented below the fish. Intuitively, the upper visual field seems to contain more relevant cues for orienting one-self. To achieve a panoramic presentation of visual stimuli covering the upper visual field, we built a compact projection setup composed of a single projector and multiple mirrors [14, 15] (Figure 1b). Larvae embedded in agarose observed view-corrected virtual 3D scenes projected onto the planar screens on the three sides, which covered azimuth of 270° and elevation of 90° (See Methods for details). The neural activity was optically monitored from above with a two-photon microscope.

**Figure 1:**
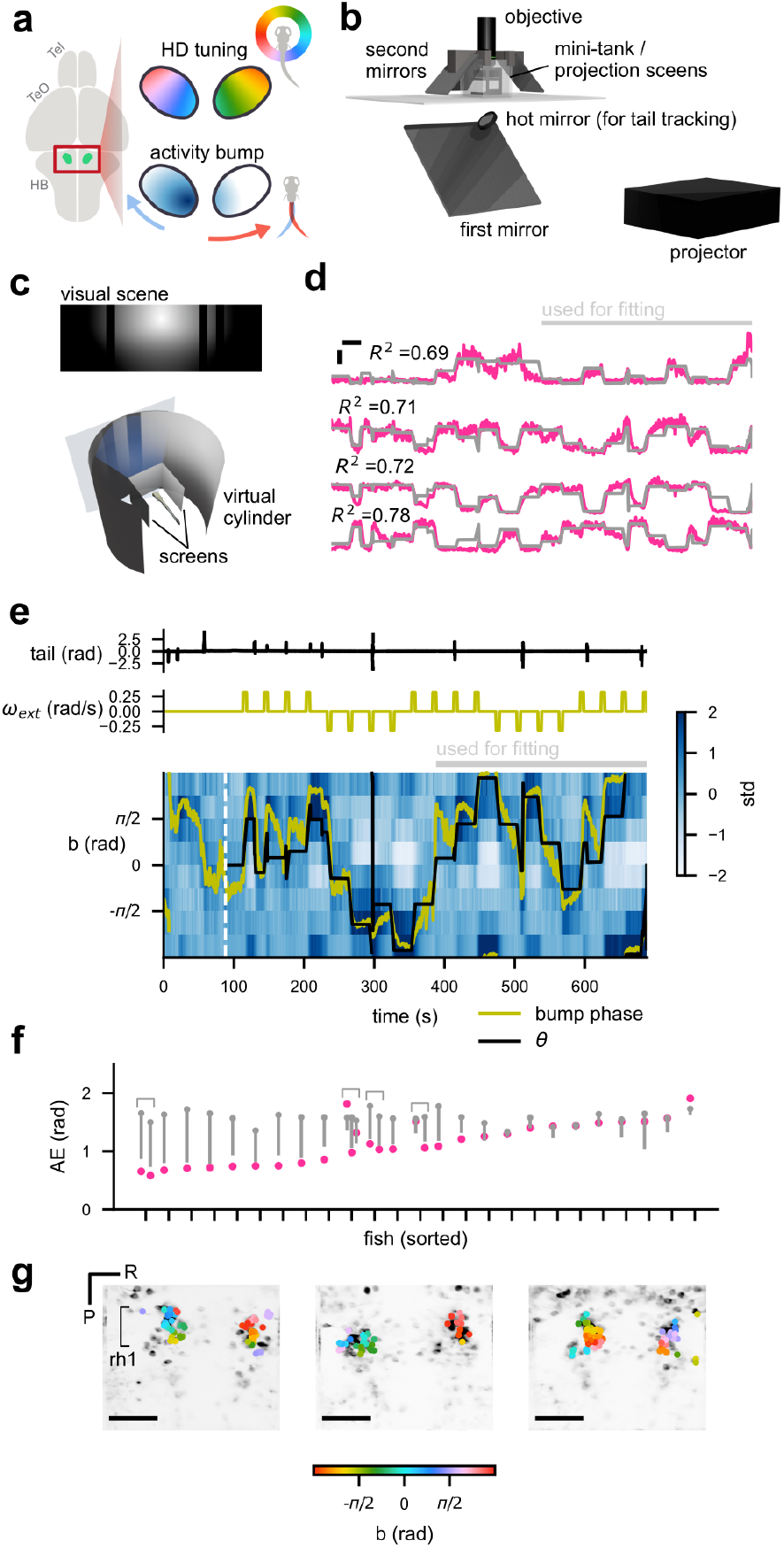
GABAergic cells in the anterior hindbrain tracks a visual scene. **(a)** The hindbrain rhombomere 1 of larval zebrafish has a cluster of gad1b+ HD neurons, whose soma locations are topographically organized according to their head direction tunings. As a result, the population activity of these cells appear as a single “bump” that move as fish turns. **(b)** Schematic of the setup. The fish was embedded in a small petri dish, and observed scenes projected around them. **(c)** The “sun-and-bars” scene consisting of a radial luminance gradient and dark bars (top) wrapped around the fish, forming a virtual cylinder (bottom). **(d)** Example activity of the cells tuned to different scene orientations (pink) with the sinusoidal fit (gray). Only the second half of the data were used for the fitting procedure. The horizontal and vertical bars respectively indicate 20 s and 1 standard deviation. **(e)** Binned activity of the scene-orientation tuned cells, with the scene orientation θ (black) and the bump phase (yellow) overlaid, as well as associated traces of the tail angle (top) as well as the exogenous rotation velocity ω_ext_ (middle). **(f)** Time-averaged absolute error (AE) (i.e., absolute differences between the scene orientation and the bump phase) for each fish (pink), compared to the shuffle (gray). The grey dots and bars respectively indicate the median and 5-percentile of the shuffle distributions. 15 out of 25 fish showed significantly below chance AE. Note that recordings from multiple planes were made in four fish, as indicated by the brackets. **(g)** Selected ROIs visualized on the anatomy, with their scene orientation tuning color-coded. Notice how the orientation preference is topographically ordered in a circular fashion in each fish, but with idiosyncratic offsets across fish. The horizontal scale bars indicate 50 µm. Tel: telencephalon; TeO: optic tectum; HB: hindbrain; std: standard deviation; R: right; P: posterior.

In this setup, we imaged the neural activity of aHB neurons expressing GCaMP6s [16] under the control of gad1b:Gal4 [17] in 6 to 9 days post-fertilization (dpf) larvae. Each experiment started with an alternating presentation of 8 s full-screen flashes of white and black. Naively visual neurons that reliably responded to these flashes were removed from the further analysis (see Methods). In the first experiment, we presented a scene consisting of a circular luminance gradient centered above the horizon and dark vertical bars (henceforth “Sun-and-bars”) for 10 minutes (Figure 1c). The scene wrapped around the fish, forming a virtual cylinder. The orientation of the scene was controlled in a closed-loop fashion based on the tail movements monitored online with a high-speed camera. In addition, slow exogenous rotation of the scene was superimposed intermittently (90° rotations over 5 seconds every 30 seconds, switching directions every 4 times), such that the fish would experience various scene orientations even if they did not make many spontaneous turns. If the HD cells indeed exploit visual information as hypothesized, their activity should be well fit by a single-peaked periodic function of the scene orientation. Thus, we fit a scaled, shifted sinusoid *a* cos (*θ − b*) + *c* to a half of the normalized fluorescence time trace of each cell. Here, *θ* denotes the orientation of the scene relative to the fish (clockwise positive), and a, b, c respectively represents response amplitude, preferred orientation, and baseline. We selected cells well fit by this sinusoid (*R*^2^ *>* 0.15) for further analyses. In this notation, the fish’s heading in the virtual world would be *−θ*.

The sinusoidal fitting identified cells tuned to the scene orientation in a majority of fish we recorded from. In some cells, the goodness of fit *R*^2^ was as high as 0.8 (Figure 1d, Figure S1a). At the population level, a single “bump” of activity was clearly visible in most fish, often even before the onset of the Sun-and-bars scene (i. e., during flash presentations) (Figure 1e, Figure S1b). We then calculated the “bump phase” as the angle of activity-weighted average of preferred orientation vectors of the selected cells [11] (see Methods). In most fish, the bump phase followed the scene orientation well, even for the period not used for the sinusoidal fitting (Figure 1f). Additionally, the preferred scene orientations of individual cells exhibited a clockwise topographical organization in rh1 (Figure 1g, Figure S1c), consistent with the previous observed topography of HD cells [7]. Interestingly, the anatomical arrangement of preferred orientations had idiosyncratic offsets across multiple fish: For example, cells preferring *θ* = 0 can be on the left or right side of the brain. This observation raises a possibility that anchoring of the HD cells to visual landmarks is not hard-wired, a point we will return to later.

### The scene orientation-tuned cells integrate turning commands

While we suspect that these scene orientation-tuned cells are the previously reported HD cells, the sinusoidal fitting procedure could also pick up visual neurons tuned to local features. To gain more confidence in their identity as the HD cells, we next asked if their bump phase move as fish makes turns even without visual feedback, as expected of HD cells. In the first experiment above, we continued recordings for another 10 minutes in the darkness after turning off the visual scene. Although we made anecdotal observations (Figure S2) where the bump phase moved as fish turned, the small numbers of swim bouts in the darkness made the quantification difficult. To facilitate fish to turn frequently without giving them visual cues for rotation, we decided to exploit the optomotor response (OMR) [18] (Figure S3a). The first half of the new experiment was mostly identical to the one in Figure 1, with a closed-loop panoramic scene with exogenous rotations. We performed the sinusoidal fitting on this half of the data to identify scene orientation tuned cells. During the second half, an array of white dots on black background translating sideways (in the virtual 3D space) was presented intermittently, facilitating fish to make turns. Once fish made a swim bout, the dots immediately disappeared without providing any rotational feedback. We then asked how the bump phase moved after each swim bout.

Figure S3b shows the data from an example fish. The bump moved clockwise and counterclockwise as fish turned left and right, as expected from the angular path integration and consistent with the previous observation [7] (Figure S3c, d). The negative correlation between the turn amplitude and bump phase shift was statistically significant across the population (Figure S3e). Overall, the results here suggest that the scene orientation tuned cells detected with the sinusoidal fits can integrate the efference copy of turning commands, supporting their identity with the previously reported HD cell population.

### The same set of HD cells can track multiple scenes

So far, we have focused on how the HD cells operate in a single visual context. In the next experiment (Figure 2), we asked if the same set of HD cells can consistently represent heading direction in multiple different visual scenes. Here, the same Sun-and-bar scene was presented in a closed-loop with the exogenous rotations. After 8 minutes, the scene was switched to what we named the “Stonehenge” scene, consisting of multiple irregularly spaced bright vertical bars over a dark background, while maintaining the same exogenous rotation sequence (Figure 2a). We identified HD cells by performing the sinusoidal fits on the first half of the data (i.e., Sun-and-bars), and asked how the bump phase behaved in the Stonehenge scene.

**Figure 2:**
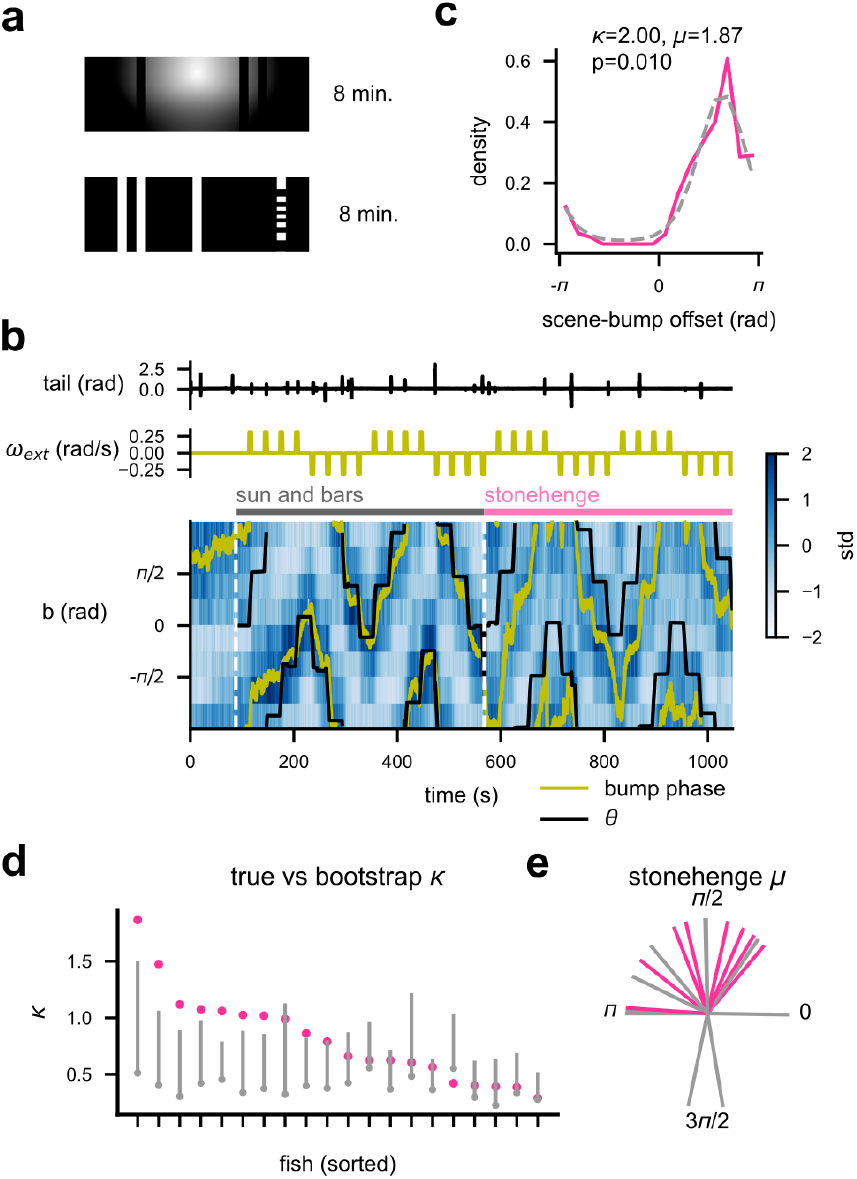
The HD neurons can track multiple scenes. **(a)** The Sun-and-bars scene (top) and the Stonehenge scene (bottom) were presented in sequence. The HD cells were identified with sinusoidal fitting on the data during the Sun-and-barsscene presentation. **(b)** Thedata from an example fish, showing the (top) tail angle, (middle) exogenousrotation velocity ω_ext_, and (bottom) the binned HD cell activity with the scene orientation θ (black) and the bump phase (yellow), as in Figure 1e. Note how the bump phase is following θ with an offset, during the Stonehenge epoch. **(c)** Histogram of the offset between θ and the bump phase during the Stonehenge epoch (pink), with a von Mises distribution fit to the data (grey dotted), which is proportional to . The p-value represents the probability that the κ (representing the peakiness of the distribution) is greater than the shuffle (see Methods for the shuffle procedure). **(d)** κ from the von Mises distributions fit on the scenebump offset histogram for each fish (pink), with the associated shuffle distribution (grey dot for the median, the bar for the 95-percentile). 8 out of 20 fish showed significantly above chance κ. **(e)** The mean angle (µ) of the von Mises distribution fit on the scene-to-bump offset during the Stonehenge epoch. Pink lines represent the data from fish with significant κ. Note how µ is not stereotypical, but is not distributed evenly from 0 to 2π.

The bump phase of the HD cells identified based on the Sun-and-bars scene significantly tracked the Stonehenge scene in about a third of the fish examined (Figure 2b-d, Figure S4), as quantified by the peakiness of the scene-bump offset distributions (*κ* of von Mises distribution fit; see Methods for details). The mean offset between the scene orientation and the bump offset was not stereotypical across fish, albeit with a bias likely due to the cross correlation between the scenes (Figure 2e). The results here demonstrate that the HD cells have a capacity to maintain stable head direction representations in various visual scenes.

### The HD cells exploit both visual landmarks and rotational optic flow

Vision can inform animals of their current head direction (e.g. distal landmarks) as well as its changes (e.g. rotational optic flow). Different circuit mechanisms are required to incorporate the two types of visual cues into the head direction estimates on a ring attractor [19]. Thus, in the next experiment, we attempted to dissociate the contributions ofthe landmark and opticflow cues on the zebrafish HD neurons (Figure 3a). The experiment consisted of three epochs. In the first (“Smooth”) epoch, the Sun-and-bars scene was presented in a closed-loop with superimposed slow exogenous rotations, similar to the earlier experiments (Figure 1, Figure 2). HD cells were identified by performing the sinusoidal fitting on this portion of data. Here, the fish had access to both landmark and optic flow cues. In the second (“Jump”) epoch, exogenous rotations were substituted by abrupt jumps of 90° without any intervening smooth rotation, eliminating the optic flow cues. If the HD cells exclusively rely on rotational optic flow to follow visual scenes, every abrupt jump should introduce a 90° discrepancy between the bump phase and the scene orientation. In the final (“Noise”) epoch, the visual scene was swapped with a featureless binary white noise with the smooth rotation sequence identical to the first epoch. If the cells relied exclusively on the landmark information to keep track of the scene orientation, exogenous rotation of the binary noise should not affect the bump phase.

**Figure 3:**
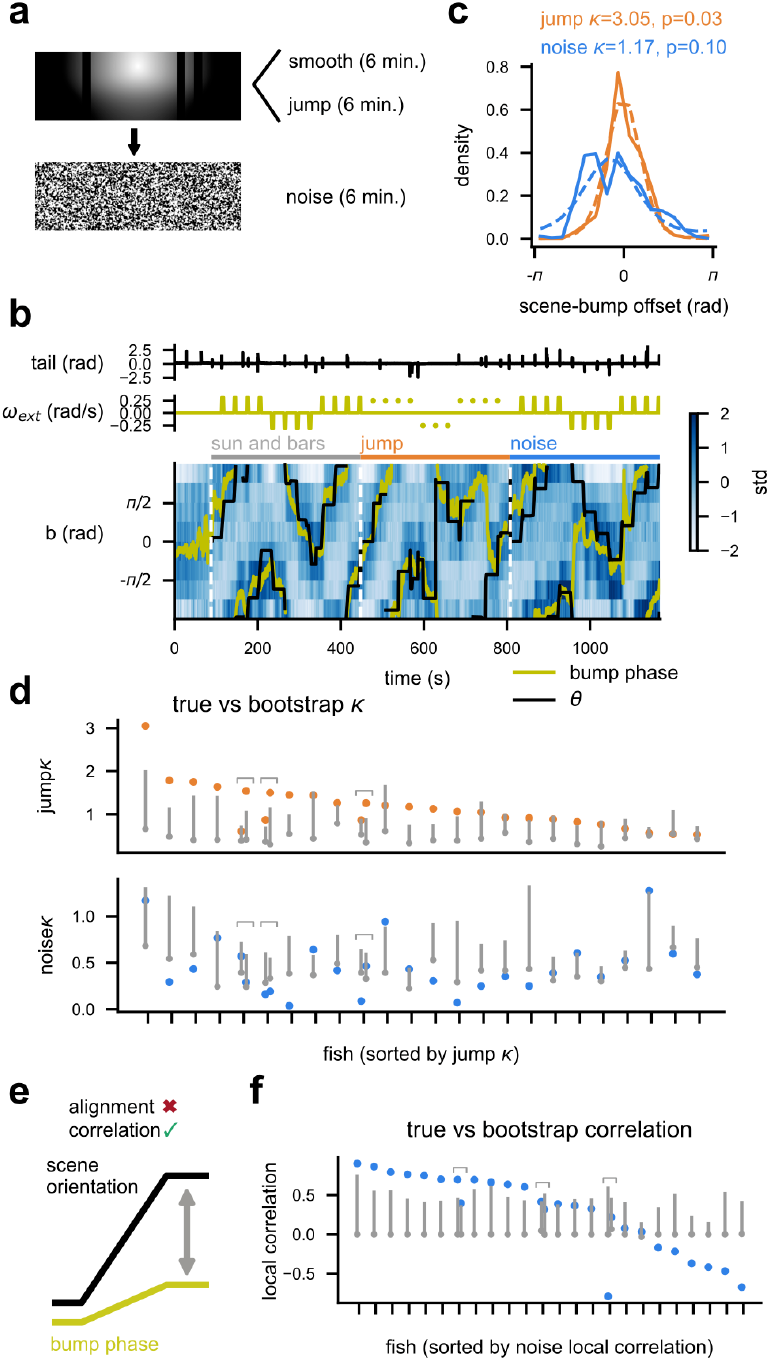
The HD neurons can exploit both landmark and optic flow cues. **(a)** Schematic of the experiment with the Jump and Noise epochs. **(b)** The data from an example single fish, showing the (top) tail angle, (middle) exogenous rotation velocity ω_ext_, and (bottom) the binned HD cell activity with the scene orientation θ (black) and the bump phase (yellow). Abrupt jumps of the scene are indicated by the dots in the middle plot. **(c)** The histograms of the scene orientation-to- bump phase offset during the Jump (orange solid) and Noise (blue solid) epochs, with von Mises distributions fit to them (dotted lines). κ of the von Mises distributions, representing the peakiness, is noted. The p-values are the probability that the κ exceeded the shuffle. **(d)** κ from the von Mises distributions fit on the scene-bump offset histogram for each fish for each condition (orange dots for Jump, blue dots for Noise). The shuffle distributions are indicated in grey (dot for the median, thebarforthe 95-percentile). Forthe Jumpepoch, 14 outof 24 fish showed significantly above change κ, whereas only 3 fish did so for the Noise epoch. Note that multiple recordings were made in 3 fish, as indicated by brackets. **(e)** If rotational optic flow moves the bump in the right direction but by an incorrect amount, the bump phase and the Noise scene orientation will be positively correlated but the offsets between the two will be variable. **(f)** Local correlation (i.e., Median Pearson correlation between θ and bump phase calculated within 15 s-windows around the exogenous rotations) in the Noise epoch (blue), compared with the bootstrap distribution (grey dot: median, grey bar: 95-percentile). 12 fish out of 24 showed significantly above chance positive correlation.

Figure 3b shows an example fish whose HD cell bump appeared to track the scene orientation well in both Jump and Noise epochs. In a majority of fish, the bump phase remained significantly aligned to the scene orientation in the Jump epoch, as quantified by the peakiness (i.e., von Mises *κ*) of the scene-bump offset distribution (Figure 3c,d, Figure S5). In contrast, only a few fish managed to maintain a constant offset between the bump phase and the scene orientation in the Noise epoch (Figure 3c,d). Still, it is possible that the rotational optic flow moved the bump in the correct direction but not necessarily for the correct amount, making the bump-scene offsets variable (Figure 3e). Indeed, in about a half of the fish, the bump phase and the scene orientation was significantly positively correlated within short (15 s) period of time around the exogenous rotation of the Noise scene (Figure 3f, Figure S5b,c) (see Methods). Overall, these results indicate that the aHB HD neurons in larval zebrafish can exploit both landmark and optic flow cues.

### Experience can alter the mapping of visual scenes to the HD cells

The idiosyncratic mapping between the soma locations of the HD cells and their scene orientation tuning (Figure 1g, Figure S1c) implied that they may be experience dependent. To observe such experience-dependency more directly, we attempted to induce remapping in the bump phase to scene orientation alignment by introducing a symmetry in the scene [20] (Figure 4a). Here, fish were again presented with visual scenes controlled in a closed-loop, with the intermittent exogenous rotations. The experiment consisted of three epochs: In the first (pre-learning) and last (post-learning) epochs, fish observed the scene consisting of a single radial gradient of luminance in the upper visual field, mimicking the sun. During the second (learning) epoch, the second sun was added to the scene azimuthally 180° away from the first one, introducing a two-fold point symmetry. In the symmetric scene, the view of a sun at a particular angular bearing is compatible with the two opposite scene orientations. If the HD cells learn this two-fold mapping, the ambiguity in the correspondence between the bump phase and the scene orientation might persist even after the removal of the second sun.

**Figure 4:**
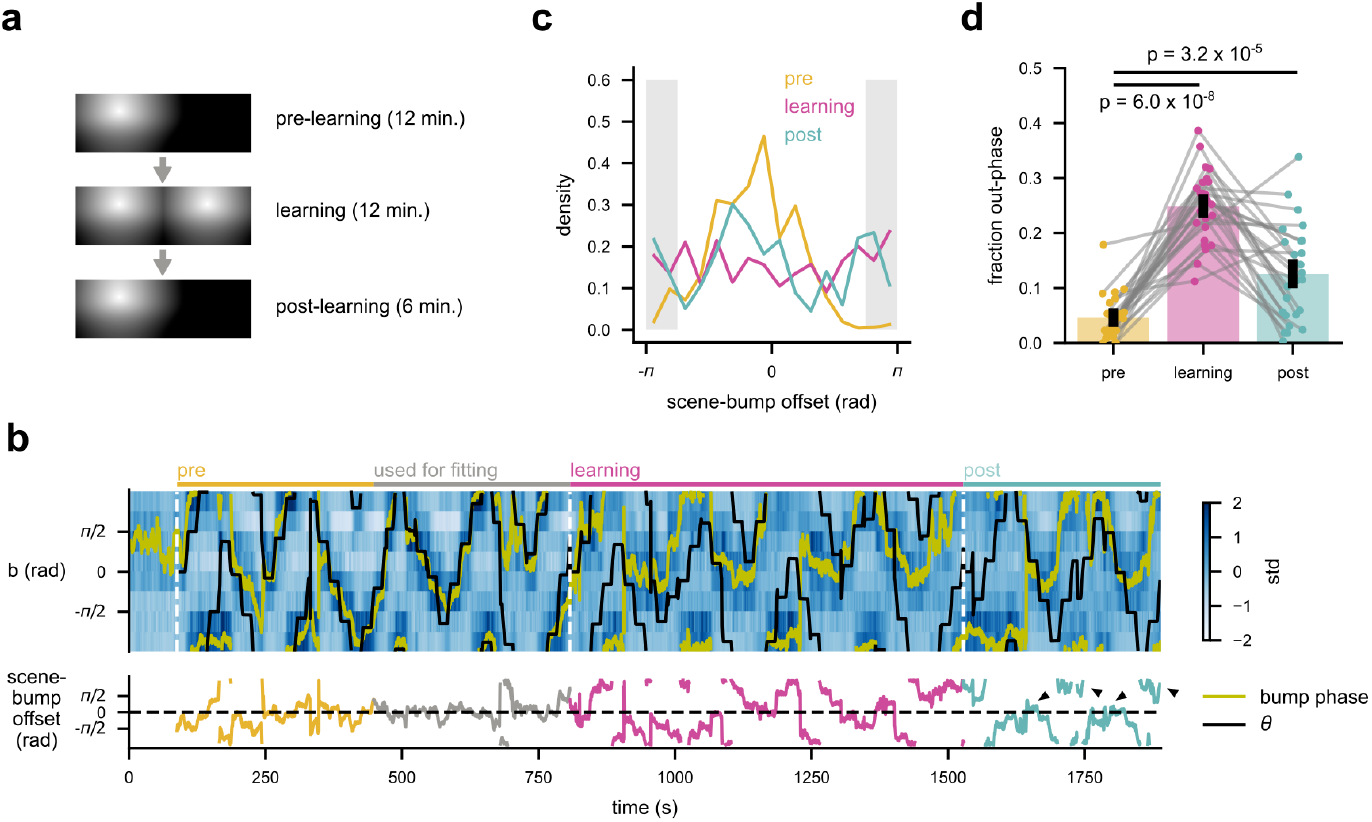
Experience-dependent remapping between the HD neuron bump and the scene orientation. **(a)** Schematic of the experiment. **(b)** Data from an example fish. (top) The binned activity of the HD cells, with the scene orientation θ (black) and the bump phase (yellow) overlaid. The HD cells were detected using the second half of the pre-learning (single sun) epoch. (bottom) The offset between the scene orientation and the bump phase over time. Places where the offset alternates between in-phase and out-phase configurations during the post-learning epochare indicated with black allows. **(c)** The histograms of the scene-bump offset for each epoch, from the fish shown in (b). The gray shaded areas indicate the out of phase range 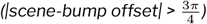. **(d)** The fraction of time that the scene-bump offset spent in the out- phase range for each epoch. The dots indicate individual fish and grey lines connect the same fish. P-values are from the sign-rank tests. N = 25 fish.

Here, we identified HD cells with the sinusoidal fitting using a half of the pre-training data. Figure 4b shows the data from an example fish. During the pre-learning epoch, the bump phase and the scene orientation generally align well, and the offset between the two shows a clear single peaked distribution centered at 0 (Figure 4c). During the learning epoch, the scene-bump offset becomes variable due to the scene symmetry (Figure 4b, c), even though the bump remained single (Figure 4b, Figure S6a). In the post-learning epoch, the bump phase appeared to deviate more from the scene, alternating between the in-phase and anti-phase configurations (Figure 4b, c). Indeed, the amount of time that the bump phase and the scene orientation were more than 135° apart significantly increased from pre-to post-learning epochs across the population (Figure 4d). A similar increase was not observed in the control group that experienced only the single sun scene throughout (Figure S6b-e). Overall, these results suggest that the anchoring of the HD cell bump to the scene orientation can be altered in an experience dependent manner.

### Inputs from the visual habenula is necessary for tracking landmarks

Finally, we asked how the visual landmark information is delivered to the HD neurons. A major source of the excitatory inputs to the dIPN, where the HD cell processes innervate, is a highly conserved epithalamic nucleus called the habenula [21] (Figure 5a). In particular, the dIPN receives asymmetric projections from the left dorsal habenula [22], which are light responsive [23] and necessary for phototaxis [24] in the larval zebrafish. Recently, we found that the left dorsal habenula neurons have azimuthally local receptive fields, and in principle capable of encoding the scene orientation as a population [25]. Thus, we hypothesized that the HD neurons receive the visual landmark information from the left dorsal habenula cells at dIPN.

**Figure 5:**
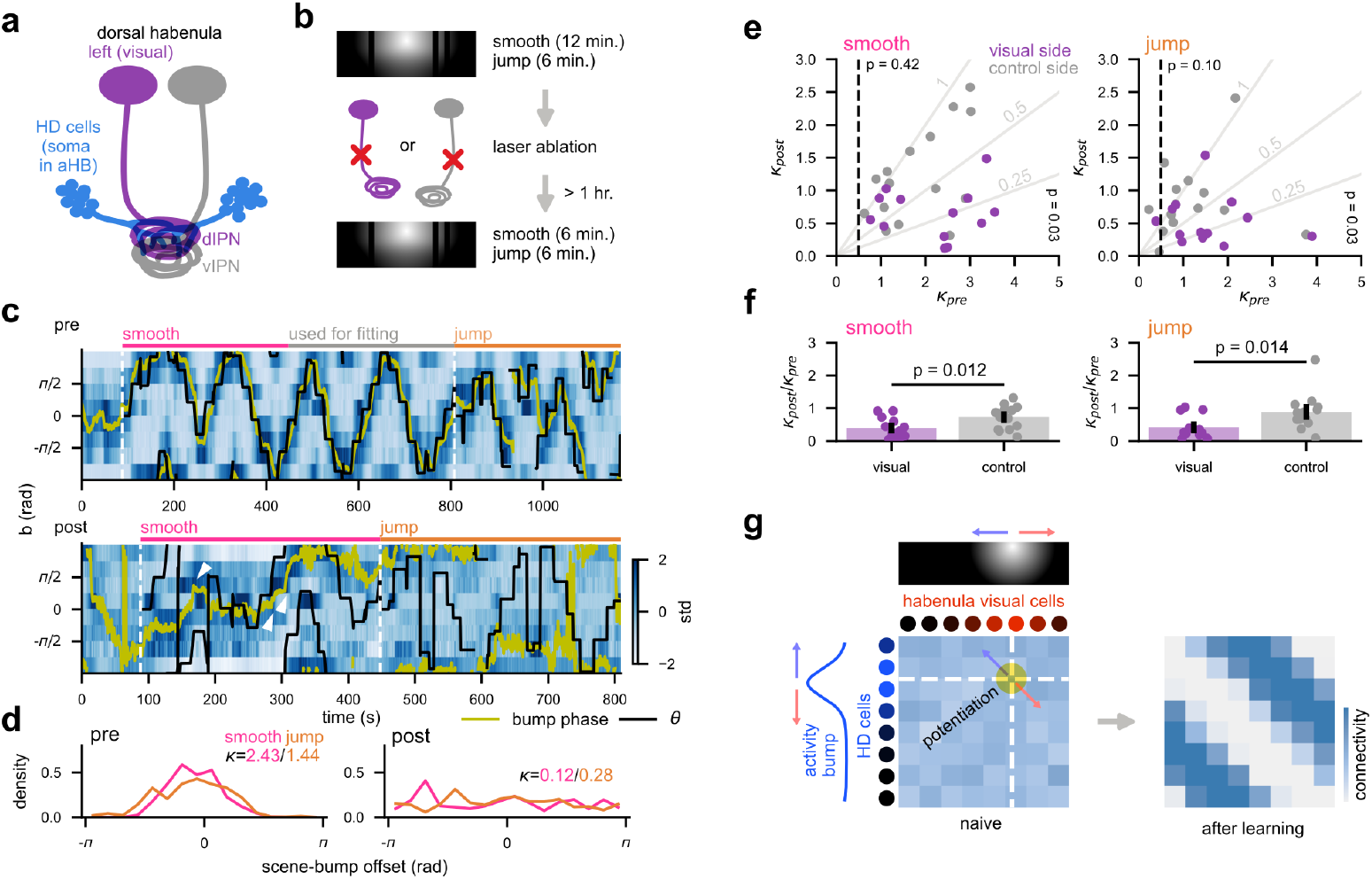
Ablating the visual habenula abolishes the tracking of the visual landmarks by the HD neurons. **(a)** Schematic of the habenulo-interpeduncular projection. Left (purple) and right (grey) dorsal habenula neuronsrespectively project to the dorsal and ventral parts of the interpeduncular nucleus (IPN). Visually responsive neurons are usually lateralized to the left habenula, and the processes of anterior hindbrain (aHB) HD neurons in nervate the dorsal IPN. **(b)** Schematic of the experiment. We repeated the presentation of the Sun-and-bars scene before and after the unilateral ablation of the habenula axons. The scene rotated smoothly about the fish, or abruptly jumped. The post-ablation recording was performed at least one hour after the ablation procedure. **(c)** The binned activity of the HD neurons before and after ablating the axons of the visual habenula, from an example fish. Note how the bump phase (yellow) follows the scene orientation θ (black) well in the pre-ablation recording (top) but not afterwards. Places where the bump phase still seemed to move in the directions of the scene rotation in the post-ablation recording are marked with white arrows. **(d)** The histograms of the scene-bump offset for pre- (left) and post-ablation (right) recordings, separately for smooth (pink) and jump (orange) epochs. κ from von Mises distributions fit on each histogram are annotated in corresponding colors. **(e)** κ of the von Mises distributions fit on the scene-bump offset histogram from the pre- and post-ablation recordings, plotted against each other. The data from Smooth (left) and Jump (right) epochs are visualized separately. Each dot represents an individual fish, and purple and grey dots respectively indicate visual- andcontrol-side ablated animals. The thin grey lines indicate different ratios between the pre- and post-ablation κ. The dotted vertical lines indicate pre-ablation κ = 0.5, and fish did not reach this threshold were excluded when comparing pre- and post-ablation κ. The p-values are from rank-sum tests performed between the two ablation groups on each axes separately. N = 13/15 fish (visual side/control) for the smooth epoch, and N = 12/13 (visual side/control) for the jump epoch. **(f)** The ratio of von Mises κ between pre- and post-ablation recordings, for each epoch type and ablation group. P-values are from rank sum tests. N = 13/15 fish (visual side/control) for the smooth epoch, and N = 12/13 (visual side/control) for the jump epoch. **(g)** Schematic of the hypothetical mechanism of how habenula inputs can anchor the HD cell bump position to visual landmarks. Habenula neurons respond to local visual features, and each of them contact the all HD neurons. As the animal explore different head directions, the activity of a habenula cell coincides more with the HD neuron tuned to specific heading that would bring visual landmarks into its receptive field. Hebbian plasticity between the two would then establish a diagonal pattern of connectivity, which makes visual landmarks instructive of the HD bump phase.

To test this hypothesis, we performed unilateral laser ablations of the habenula axons entering the IPN (see Methods for details) (Figure S7a). Before and after the ablation, we repeated an experiment similar to Figure 3, where the Sun-and-barsscene waspresented in a closed loop, with smooth rotations or abrupt jumps superimposed intermittently (Figure 5b). We also recorded responses of the habenula neurons to full-field flash stimuli to account for the inversion of the habenula laterality that happens in a minority of fish [26] (Figure S7b, c). In most fish, the population-wide correlation structure of the HD cells was maintained before and after the ablation (Figure S7d-f), suggesting that we managed to successfully identify cells across recordings. In the example fish shown in Figure 5c, d, the ablation of the visual habenula axons abolished the alignment between the scene orientation and the bump phase in both epochs, even though the bump of activity itself is still clearly visible [25]. Such decrease was not observed in the control side-ablated fish shown in Figure S7g. At the population level, the fish in the visual side ablation group exhibited lower peakiness of the scene-bump offset distribution in both epochs, only after the ablation procedure

(Figure 5e). In addition, the magnitude of the decrease in the bump-scene alignment was significantly higher in the visual side ablated group compared to the control side ablated group (Figure 5f). Interestingly, we made several anecdotal observations in the visual side ablated fish where the bump appeared to follow the smooth rotation of the scene despite the overall loss of the bump-scene alignment (Figure 5d, Figure S8). Together, these results establish that the habenular input to the dIPN is necessary for incorporating the landmark information to the HD representation, but likely not for integrating rotational optic flow.

## Discussion

The present study demonstrated that the larval zebrafish HD neurons incorporate both allothetic (i.e., landmarks) and idiothetic (i.e., opticflow) visual cuesinto their heading estimate. In particular, we found that the plastic mapping between the scene orientation and the bump phase requires inputs from the visually responsive side of the habenula to dIPN. How does the microcircuitry in the dIPN implement the mapping between the heading and visual landmarks? A plausible mechanism is the Hebbian plasticity between the habenula axons and the HD neurons, similar to those found between the Drosophila ring (R) neurons and HD neurons (i.e., E-PG cells) [20, 27]. In fact, morpho-physiological parallels between the habenula neurons and the fly R cells are rather remarkable: they both have local visual receptive fields [25, 28], and their individual axons wrap around the entire lateral extent of the output neuropil [29, 30], contacting the whole suite of HD neurons [31, 32]. Here, the synaptic weights from a habenula neuron to the HD cells are assumed to be initially random. As the animal explores the environment, the habenula cell activity would coincide frequently with the activity of HD cells tuned for the specific heading that brings a reliable distal landmark (e.g., the sun) into the receptive field of the habenula cell, strengthening their connectivity (Figure 5g). At the population level, this process would establish topographically ordered connectivity from the habenula to the HD cells with arbitrary offsets. Once such connectivity is established, the visual landmark becomes instructive to the HD cell activity. While our data is also compatible with the possibility that the visual landmark information arrives at the dIPN from elsewhere and the habenula inputs only modulate the plasticity, we find this unlikely, given that (1) neuromodulatory neurons in the habenula are lateralized to the non-visual side [33], and that (2) no other obvious visual inputs to the dIPN are known . In addition to visual landmarks, the habenula responds to diverse sensory modalities, such as translational optic flow [25], odor [23], and temperature [34], again similar to how the fly R neurons are multimodal [35]. This raises the possibility that the HD neuron activity can be tethered to other types of allocentric cues with similar mechanisms of plasticity.

Unlike the landmark information, it remains less clear how the idiothetic information arrives at the dIPN and how the HD neurons integrate it. Ring attractor models typically implement the angular path integration by assuming groups of cells that asymmetrically connect neighboring HD neurons and become active only when the animal makes a turn in a particular direction [3]. These turn-gated asymmetric connections would move the bump phase in appropriate directions. In Drosophila, neurons called P-EN asymmetrically connect the HD cells (i.e., E-PG) [36], and are modulated by multimodal cells called GLNO, which construct multi-modal estimates of the animals’ self-motion combining the motor efference copy and optic flow signals [37]. In mammals, the nucleus prepositus hypoglossi (NPH) of the rostral medulla is suspected to be informing the HD cells of the angular head velocity. The NPH receives inputs from vestibular nuclei and projects to the DTN [38, 39], and is necessary for accurate path integration in the thalamic HD cells [40, 41]. While the larval zebrafish homolog of the NPH is unknown, several clusters of cells tuned to rotational optic flow have been identified in the larval zebrafish medulla [42–44]. It is of particular future interest to check if these medulla cells send homologous projections to DTN, and how they contribute to the angular path integration in the dIPN.

In mammals, the three major brain regions containing the HD neurons are the lateral mammillary nucleus (LMN), the anterodorsal nucleus of the thalamus (ADN), and the postsubiculum (PoS), which form a loop and maintains a coherent representation of heading [13, 45]. The anchoring of the heading representation to visual landmarks are thought to happen through interactions between the retrosplenial cortex and PoS [46, 47], which is then propagated back to the subcortical areas through feedback projections [48]. In contrast, the observations of HD cells in the teleost forebrain are limited [49], and overall the homology of mammalian thalamo-cortical connections in fish is questioned [50, 51]. In light of the highly conserved nature of the habenula-IPN projection [21], our results suggest an evolutionary scenario where the habenula-IPN mechanisms for the landmark anchoring similar to those of insects first evolved in early vertebrates, and the forebrain circuitry was added on top of them, as the elaborate visual telencephalon evolved in mammals.

## Acknowledgements

The authors thank the members of the Portugues lab for constructive discussions on the project. We want to thank Martin Haesemeyer for sending us the Tg(18107:Gal4) line. R. T. was supported by the European Molecular Biology Organization (EMBO ALTF 732-2022) as well as the Human Frontier Science Program (HFSP LT0027/2023-L) for this work. This research was funded by the German Research Foundation (DFG) under Germany’s Excellence Strategy within the framework of the Munich Cluster for Systems Neurology (EXC 2145 SyNergy, identifier 390857198), through the “Enhanced resolution microscopy” project DFG – Projekt-nummer 518284373, by the Volkswagen Stiftung via a Life? grant and by the Max Planck Gesellschaft.

## Materials and Methods

### Zebrafish husbandry

The animal handling and experiments were performed according to protocols approved by the Technische Universität München (TUM) and the Regierung von Oberbayern (animal protocol number 55-2-1-54-2532-101-12 and 55.2-2532.Vet_02-24-5). Adult zebrafish (Danio rerio) were housed in the facility at the Institute for Neuronal Cell Biology at TUM. The adult fish were maintained in water temperature of 27.5 – 28.0 °C on the 14:10 hour light:dark cycle. All experiments were performed on 6 to 9 days post fertilization (d. p. f.) larvae of undetermined sex. The eggs were kept in 0.3x Danieau solution, and in the water from the fish facility upon hatching. The larvae were maintained at 28.0 °C and under the 14:10 hour light:dark cycle.

### Animal strains

All imaging experiments were performed on fish carrying Tg(gad1b:Gal4)mpn155 [17] and Tg(UAS:GCaMP6s)mpn101 [52]. To label the habenula, a previously uncharacterized enhancer trap line Tg(18107:Gal4) was used. The expression pattern of the 18107:Gal4 can be browed as Z Brain Atlas (https://zebrafishexplorer.zib.de/home/). A subset of fish with the habenula labeling possessed Tg(UAS:nfsB-mCherry) [53] for a logistical reason. This does not affect the results of the ablation experiments as they were evenly distributed across the conditions, and the fish were not treated with relevant chemogenetic reagents. All fish were mitfa -/-(i.e., nacre) mutants lacking melanophores to allow optical access to the brain.

## Two-photon microscopy experiments

### Animal preparation and the stimulus presentation setup

Animals were embedded in 2% low-melting point agarose in 30 mm petri dishes. The agarose around the tail was carefully removed with a scalpel to allow tail movements. The dish was mounted on a 3D-printed pedestal and placed in a cube-shaped acrylic tank with the outer edge length of 51 mm. The height of the pedestal was designed so that the head of the animal comes at the center of the tank, taking the thickness of the dish and the typical amount of agarose into account. The tank was then filled with the fish facility water to minimize the refraction due to the petri dish wall. The three sides of the tank (except for the one facing the back of the fish) were made of single-side frosted acrylic (PLEXIGLAS Satinice 0M033 SC), which functioned as projection screens. The diffusive side faced inward to minimize the reflections between the walls. The visual stimuli were projected onto the three walls through two sets of mirrors with a previously described geometry [15] (Figure 1), subtending 270° horizontally and 90° vertically. The larvae were lit with an IR LED array through the transparent back wall of the tank. Their tail movements were monitored from below with a high-speed camera (Allied Vision Pike F032) at 200 Hz, though a hot mirror and a short pass filter to reject the excitation beam.

#### Microscope

Functional imaging was performed with a custom-built two photon microscope. The excitation was provided by a femtosecond pulsed laser with the wavelength of 920 nm, the repetition rate of 80 MHz, and the average source power of 1.8 W (Spark ALCOR 920-2). The average power at the sample was approximately 10 mW. The scan head consisted of a horizontally scanning 12 kHz resonant mirror and a vertically scanning galvo mirror. Pixels were acquired at 20 MHz and averaged 8-fold, resulting in the frame rate of 5 Hz. The typical dimension ofthe image was about 100 um x 100 um, with the resolution of about 0.2 um per pixel. Only pixels corresponding to the middle 80% of the horizontal scanning range were acquired to avoid image distortion, and the area outside was not excited to minimize photo-damage. The fast power modulation was achieved with the acousto-optic modulator built in to the laser. Imaging was targeted at the anterior hindbrain region previously reported to contain gad1b+ HD neurons [7], as well as the bilateral dorsal habenula (Figure S7).

#### Stimulus protocols

All visual stimulus presentation and behavioral tracking were performed using Stytra package [54]. The panoramic virtual reality environments were created and rendered using OpenGL through a python wrapper (ModernGL). Because we were only concerned about the angle of the animal with respect to the virtual world, all the scenes were simulated as textures mapped onto a virtual cylinder surrounding the fish. The texture had the dimension of 720 x 240 pixel, thus each pixel subtended about 0.5°. Dynamic aspects of the scenes (i.e., scene rotations and movements of the dots) were achieved by updating the textures on the cylinder. In each frame of the stimuli, three views corresponding to the three screen walls were rendered, which were arranged on the projector window to fit the screens.

Behavioral tracking was performed as reported previously [54]. Briefly, seven to nine linear segments were fit to the tail of the larva, and the “tail angle” was calculated at each camera frame as the cumulative sum of the angular offsets between the neighboring pairs of segments. To detect swimming bouts, a running standard deviation of the tail angle within a 50 ms window was calculated (“vigor”). A swimming bout is defined as a contiguous period during which the vigor surpassed 0.1 rad. For each bout, the average tail angle within 70 ms after the onset was calculated, with a subtraction of the baseline angle 50 ms prior to the bout onset. This average angle (“bout bias”) captures the first cycle of the tail oscillation in a bout and correlates well with the heading change in the freely swimming larvae [55]. Once the bias is calculated, the scene was rotated by the same amount as the bias, with the velocity profile of a decaying exponential with the time constant of 50 ms.

The visual scenes used in the study were as follows:

- Flash: Uniform fields of black or white.
- Sun-and-bars: Three black vertical bars on a single radial gradient of luminance, ranging from white at the center and black at the periphery. The bars were 15° wide and respectively centered at -90°, +75°, and +105° azimuths (0° is to the front and positive angles to the right). The center of the gradient was in front and 45° above the horizon, and the radius was 135°.
- Translating Dots: Dots randomly distributed in a virtual 3D space at the density of 7.2 cm-3 moved at 10 mm/s sideways. The dots within the 40 mm cubic region around the observer were rendered as 3 x 3-pixel white squares against a black background, regardless of the distance.
- Stonehenge: Four while vertical bars on a black background. The bars were respectively positioned at -120°, -90°, 0°, and +135° azimuths. The right-most bar was broken in the vertical direction with the periodicity of 20° elevation and the duty cycle of 50%.
- Noise: A 2D array of uniform random numbers within [0, 1], smoothed with a 5 x 5 2D boxcar kernel and binarized into black and white at 0.5.
- Single-sun: A radial gradient of luminance from white at the center and black at the periphery, centered at -90° azimuth and 35° above the horizon, with the radius of 60°.
- Double-sun: The same as the Single-sun scene, but symmetrized around the vertical meridian.

In order to identify and exclude naively visual neurons, each aHB recording started with the alternating presentation of white and black flashes (8 s long each, 5 repetitions). In the experiment in Figure S3, the Translating Dots moving leftward and rightward alternatingly were also presented (8 s long, 5 repetitions). Afterwards, epochs of closed-loop scene presentations started. At the beginning of each epoch, the scene orientation was reset to 0°. On top of the closed-loop control, episodes of exogenous slow rotation (18 °/s) was superimposed intermittently (5 s every 30 s (Figures 1-3, S1-S5) or 20 s (Figures 4, 5, S6-S8)). The directions of the rotations flipped after every 4 rotation episodes. The structures of the virtual reality experiments were as follows:

- Figure 1 (S1, S2): In the first epoch (10 min.), the Sun-and-bars scene was presented. In the second epoch (10 min.), fish received no visual stimuli (i.e., darkness).
- Figure S3: In the first epoch (8 min.), the Sun-and-bars (N = 10 fish) or Stonehenge (N = 15 fish) scene was presented. In the second epoch (15 min.), the fish observed Translating Dots moving either left or right. The dots disappeared and the screen turned uniform white if the fish performed a bout or 10 seconds passed without a bout. The dots reappeared after waiting for 10 seconds.
- Figure 2 (S4): In the first and second epochs (8 min. each), the Sun-and-bars and the Stonehenge scenes were presented, respectively.
- Figure 3 (S5): In the first and second epochs (6 min. each), the Sun-and-bars scene was presented. In the second epoch, the superimposed exogenous rotations were swapped with abrupt 90° jumps. In the third epoch (6 min.), the Noise scene was presented.
- Figure 4, (S6): In the first and third epochs (12 min. each), the Single-sun scene was presented. In the second epoch (12 min.), either the Double-sun (N = 25 fish) (Figure 4) or Single-sun (N = 20 fish) scene (Figure S6) was presented.
- Figure 5 (S7, S8): In all epochs, the Sun-and bars scene was presented. The first epochs were 12 and 6 min. long in the pre- and post-ablation recordings, respectively. In the second epochs (6 min.), the superimposed exogenous rotations were swapped with abrupt 90° jumps.

In addition, we recorded the responses of the habenula neurons to alternating black and white flash stimuli (8 seconds each, 10 repetitions) to determine the visually responsive side.

#### Laser ablations

The habenula axons (i.e., fasciculus retroflexus; FR) were ablated unilaterally either by scanning a laser within a small ROI on the FR (Setup A: Spectra Physics MaiTai, 830 nm, source power 1.5 W) or by pointing a laser on the FR (Setup B: Spark ALCOR 920-2, 920 nm, source power 1.8 W). The pulsing characteristics of the two lasers were comparable (repetition rate 80 MHz, pulse width < 100 fs ), but ALCOR was group delay dispersion-corrected and thus more efficient for ablations. On the setup A, scanning with the duration of 200 ms was repeated 10 times with the interval of 300 ms. On the setup B, a couple 100 ms long pulses were delivered. These procedures were repeated until the successful ablation was confirmed either by a spot of increased fluorescence due to the damage, or a cavitation bubble. Ablations were repeated at two to three locations around the midbrain/pretectum levels to ensure the complete cut of the FR. The numbers of the fish treated on the setups A and B were respectively N = 8/8 (visual side ablated/control) and 5/7 (visual side ablated/control) fish. We waited at least one hour after the ablation before making the post-ablation recordings.

## Data analysis

### Preprocessing

All imaging data were pre-processed using the suite2p package [56]. Briefly, frames were iteratively aligned to reference frames randomly picked from the movie, using phase-correlation. To detect regions of interests (ROIs) representing cellular somata, a singular value decomposition was performed on the aligned movie, and the ROIs were seeded from the peaks of the spatial singular value vectors. All ROIs were used without morphological classifiers for cells, because functional ROI selection procedure described below rejected spurious non-cell ROIs. In the ablation experiments, ROIs were manually defined, as described below.

### ROI selection

Fluorescent time traces from the anterior hindbrain recordings were first normalized into Z-scores for each ROI by subtracting the mean and dividing by the standard deviation. The normalized traces were then smoothed with a box-car kernel with the width of 1 second. Next, a scaled shifted sinusoid *a* cos(*θ− b*) + *c* was fit to the smoothed traces of each ROI, where *θ* is the orientation of the visual scene relative to the fish. The fitting was performed with the “curve_fit” function from the scipy package, and parameters were bounded in the range of *a ≥* 0, *b ∈* (*−π, π*]. Only fractions of the data were used for this fitting to allow cross-validated quantifications of bump-to-scene alignments on the held-out fraction (described below). Specifically, either the second half of the first epoch (Figures 1, 4, 5), the entire first epoch (Figures 2, 3, S3) were used for the fitting. ROIs with *R*^2^ larger than 0.15 were considered to be sufficiently modulated by the scene orientation and included. In addition, pairwise Pearson correlations of response time traces to repeatedly presented flashes (and translating dots in Figure S3) at the beginning of the experiments were calculated for each ROI, and averaged across all pairs of repetitions. Naively visual ROIs with the mean correlations above 0.1 were excluded from the further analyses. Finally, rectangular masks were manually drawn around the rhombomere 1 of the anterior hindbrain, and the ROIs outside the mask were excluded.

### Ablation experiment-specific preprocessing

For each fish in the ablation experiment (Figure 5), we first analyzed the responses of the habenula neurons to the repeated flash stimuli. We calculated the mean correlation (as defined in the “ROI selection” section) to repeated black and white flashes, and counted the numbers of ROIs on each side that had the mean correlation > 0.3 (“visual ROIs”). The side with more visual ROIs was defined as the visual side. In a minority of fish where the number of the visual ROIs from the two sides were close (within ±1 range), we determined the visual side based on the morphology. This was possible due to the asymmetric expression of the 18107:Gal4. Overall, we found 3 fish with inverted laterality (i.e., right visual) out of 28 included in the analysis.

To compare the behaviors of the HD neurons before and after the FR ablations, ROIs were manually defined about the cells that were identifiable in both pre- and post-ablation recordings. To minimize the effort for manual ROI drawing, we first used the ROIs detected by the suite2p pipeline from the pre-ablation recording, and run the HD cell selection procedure as described above. Using this as a guide, we drew manual ROIs on the pre-ablation recording, specifically focusing on the suite2p-based HD cell ROIs. We then calculated affine transformation between the average frames of the pre- and postablation recordings, and used this transform to register the manually defined ROIs to the post-ablation recordings. Finally, we manually adjusted the ROIs to better match the average post-ablation frames, if necessary. To make sure that we managed to identify the same cells across two recordings, we calculated the Pearson correlations of the smoothed fluorescence traces between the all pairs of ROIs for each recording, and then computed the correlation of those correlations. We excluded fish with correlations of pairwise correlations below 0.4 from the further analyses.

### Bump phase calculation

As a readout of the population-level, instantaneous estimate of the scene orientation *θ*(*t*) by the HD neurons, we calculated the “bump phase” 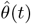, where t is discretized time. To do so, we first averaged the ROI-wise response time traces within eight 45° bins of the preferred orientation b. We excluded fish with more than 4 empty bins from the following analyses. We then computed the bump phase as

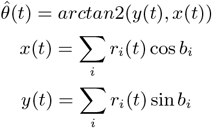

where *r*_*i*_(*t*) is the average response of the i-th bin at time *t*, and *b*_*i*_ is the central angle of the i-th bin. The bump amplitude was also calculated as 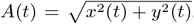. To examine if the bump phase followed the scene orientation, we calculated the centered scene-bump offset 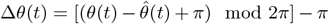 . In addition, to examine if the bump phase moved in the same directions as the scene moved (Figure 3), we calculated Pearson correlations between unwrapped *θ*(*t*) and *θ*(*t*) within 15 s windows centered about the episodes of 5 s exogenous rotations. We then calculated the median of these correlations over the all rotation episodes, which we termed “local correlation”.

Statistical quantifications

In Figure 1, we quantified the alignment between the bump phase and the scene orientation as the “absolute error” defined as *AE*_*data*_ = ∫ |Δ*θ*| *dt*, using the held out data (i.e., the first half of the first epoch). To test its significance, we performed a bootstrap test: for each fish, we circularly shifted *θ*(*t*) in time randomly, and recalculated the absolute error (*AE*_*BS*_ ) 1,000 times. We tested the significance of *AE*_*data*_ as the probability *P* (*AE*_*BS*_ *< AE*_*data*_).

In Figure S3, we calculated correlations and slopes between the biases of the bouts with the change in the bump phase for each fish. The change in the bump phase is calculated as the difference between *θ*(*t*) 5 seconds after the bout minus the average *θ*(*t*) during 1 second preceding the bout. For this calculation, we selected bouts that happened in the second epochs (i.e., darkness or intermittent translating dots) that were separated for more than 5 seconds from preceding or following bouts. A minority of bouts that had a bias >120° as well as ones where tail tracking was faulty (tail not tracked >10% of frames) were discarded as unreliable. Fish that did not have at least 5 such good bouts were excluded from the analyses. We had a single fish in the dataset that had two recordings that passed the above quality thresholds. The correlations and slopes from multiple recordings in this fish were averaged.

In Figures 2, 3, the bump-scene alignment was examined by how peaky the distribution of Δ*θ* was. To do so, we first calculated the normalized histogram of Δ*θ* with 16 equally spaced bins spanning ( *−π, π*], on which we fit von Mises probability density functions *f* (*x*) = *Ce*^(*κ* cos (*x−µ*))^, where *µ* controls the mean and *κ* the peakiness(*C* isa scalarto ensure *f* (*x*) integrates to 1). We then performed 1,000-fold bootstrap tests with time-domain shifting as above, calculating the significance of *κ* as *P* (*κ*_*BS*_>*κ*_*data*_) for each fish. This test is more liberal than the test on the absolute error in that it allows the realignment between the bump phase and the scene orientation, due to the difference in visual scenes or re-learning. The significance of the local correlation (Figure 3) between unwrapped *θ*(*t*) and *θ*(*t*) was also tested for each fish with 1,000-fold bootstrap tests with random circular shifting in the time domain. In calculating the local correlation from shifted *θ*(*t*) , the 15 s-long windows containing the discontinuous point created by circular shifting was discarded.

In Figure 4, the alignment between the bump phase and the scene orientation was quantified by the fraction of time Δ*θ* was in the anti-phase range 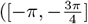 and 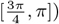, in line with a previous study with a similar behavioral protocol [20]. This anti-phase fraction was calculated separately for each of the three epochs (only using the data not used for the sinusoidal fitting) and compared across the epochs with the signed-rank tests. In addition, the difference in the anti-phase fractions between the first (pre-training) and the last (post-training) epochs were compared across the groups presented with the Double- or Single-sun scenes during the training, using the rank-sum test. For these comparisons, a minority of fish whose bump amplitude A decayed more than 60% between the pre- and post-training epoch (e.g., due to poor health) were discarded as unreliable.

In Figure 5, the peakiness of the Δ*θ* distributions were quantified by *κ* of the von Mises fits, as described above. For each fish, *κ* was separately calculated for each epoch, pre- and post-ablation recordings, only using the data not used for the sinusoidal fitting. We then compared the *κ* across the ablation group within each epoch of each recording with signed-rank tests. In addition, the ratios of *κ* between pre- and post-ablation recordings for each epoch were compared across the ablation groups, using the rank-sum test. We excluded fish that had *κ <* 0.5 in the pre-ablation epochs, as they were not informative about the effect of the ablations.

## Data and code availability

All the data described in the paper, the analysis notebooks and the figure generation notebooks will be made available upon publication.

## Supplementary figures

Eight supplementary figures follow.

**Figure S1:**
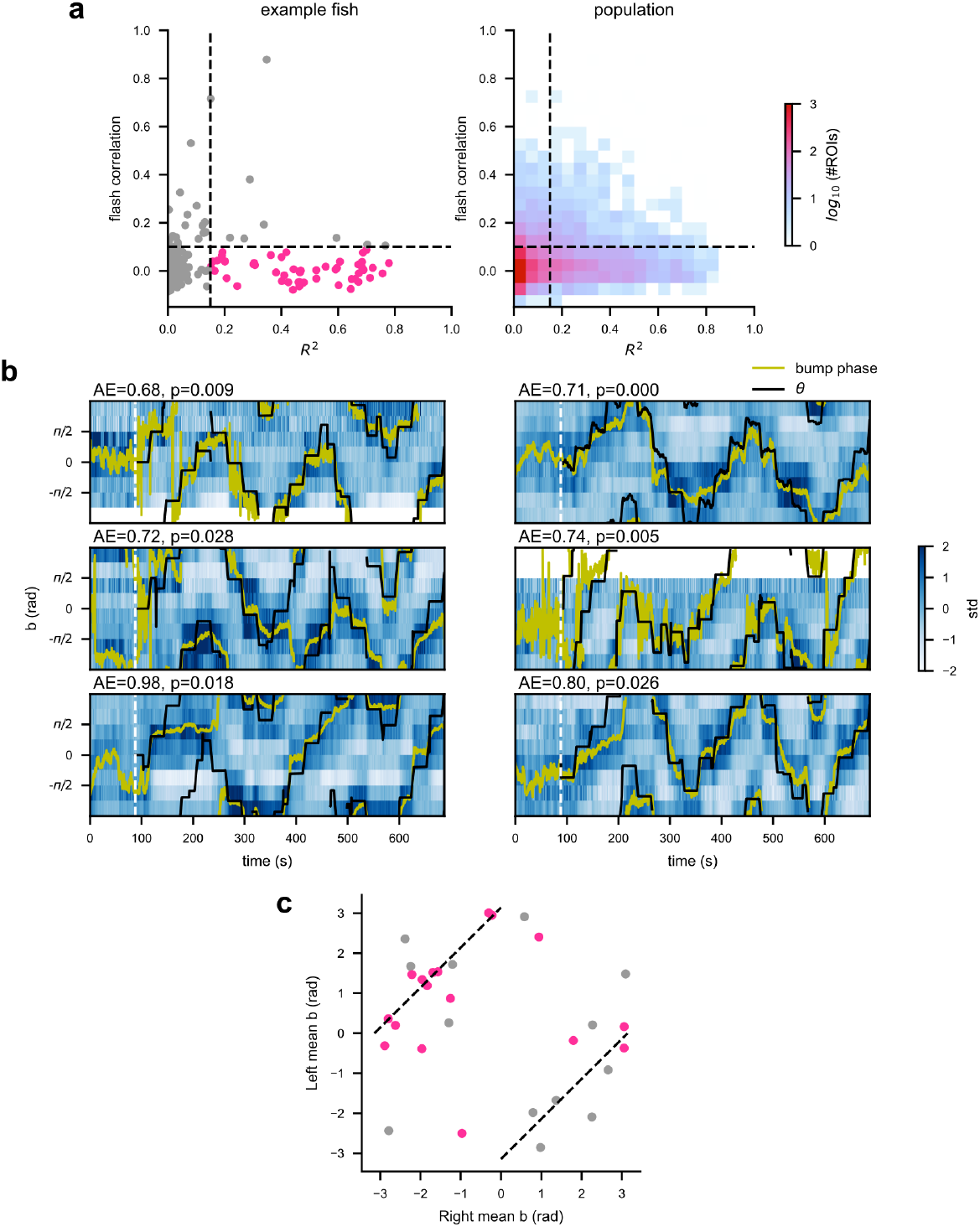
Behaviors of the scene orientation tuned cells in the Sun-and-bars scene. **(a)** The distribution of the R^2^ values from sinusoidal fits and the mean pairwise correlation to the flash stimuli, for the same fish as in Figure 1e (left) and for the population (right). **(b)** The binned activity of the scene orientation tuned cells from six more example fish as in Figure 1e. The dotted line marks the beginning of the Sun-and-bars scene. Only the second half of the epoch was used for the sinusoidal fitting. The time-averaged absolute error (AE) and associated p-values from the bootstrap tests are indicated. **(c)** Mean preferred scene orientations b of the ROIs in the left and right hemispheres, plotted against each other. Pink dots are for the recordings where the absolute error was below chance. The data roughly lay on top on the diagonal dotted lines indicating (left mean – right mean) = π. N = 30 recordings from 25 fish.

**Figure S2:**
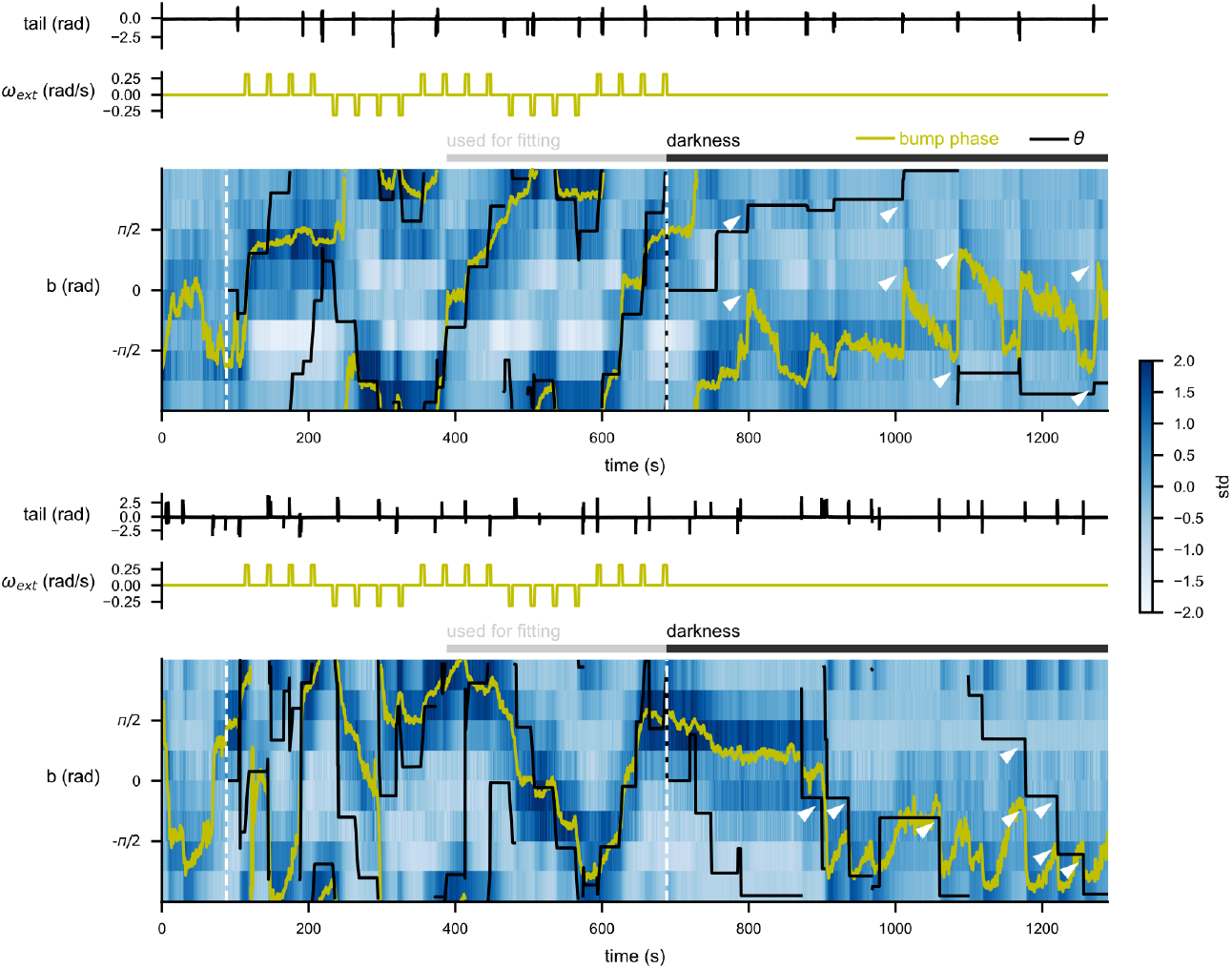
Examples of the bump phase movements in the darkness. Binned activities of the scene orientation tuned cells from two example fish throughout the entire recording, as in Figure 1e. The dotted white lines indicate the beginnings of the Sun-and-bars and darkness epochs, respectively. Places where lateralized swim bouts in the darkness appeared to coincide with bump phase changes are marked by white triangles. Interestingly, in both fish here, the bump appeared to drift in one sense and fish appeared to swim to counteract the drift.

**Figure S3:**
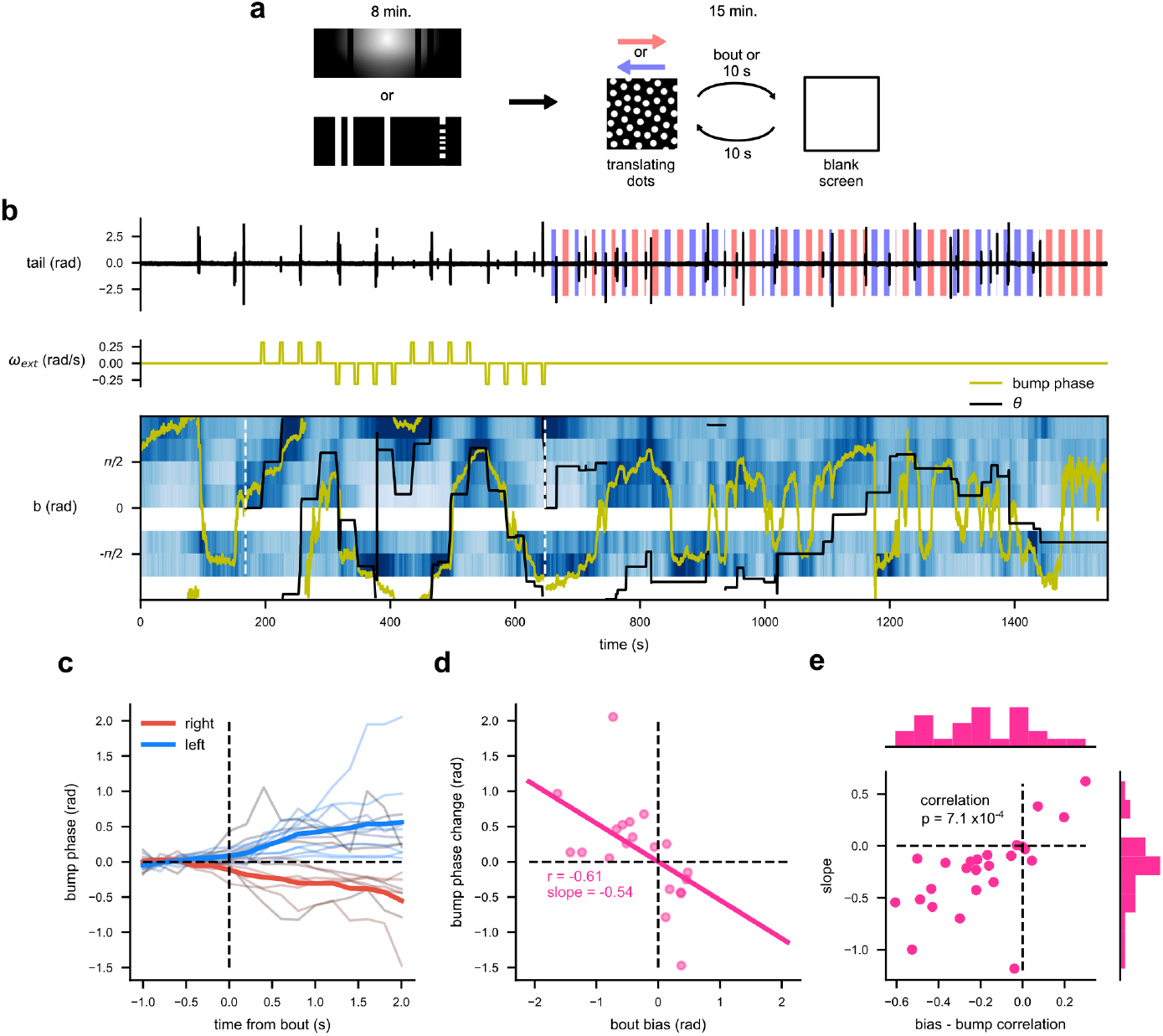
Turns triggered by translational optic flow move the bump phase. **(a)** Schematics of the experimental structure. During the first epoch, the fish experienced a panoramic scene rotating about them. During the second epoch, the fish was presented with dots translating leftward or rightward, which facilitate them to make turns. Once fish made a swim bout or 10 seconds passed without a bout, the dots were turned offand screenwent blank. The dots reappeared in another 10 seconds. **(b)** The binned activity of the scene orientation-tuned cells in a single example fish detected from the first epoch, as in Figure 1e. The calculated bump phase (yellow) and scene orientation θ (black) are overlaid (note that θ did not correspond to the stimuli presented in the second epoch). The red and blue boxes on the tail plot respectively indicate periods during which rightward and leftward dots were presented. **(c)** The changes in the bump phase around each swim bout (thin lines). Thick lines are the averages for the right and left bouts (defined as bias > 0.2 and < -0.2 rad, respectively). **(d)** The bump phase change (i.e., average bump phase between 1 to 2 seconds after the bout onset, with a baseline subtraction) plotted against the bout bias. **(e)** The correlation and slopes between the bump phase change and bout bias distributions, plotted against each other. The histograms show the marginal distributions. The bout bias-bump phase change correlation was significantly negative across the population (signed-rank test, N = 25 fish).

**Figure S4:**
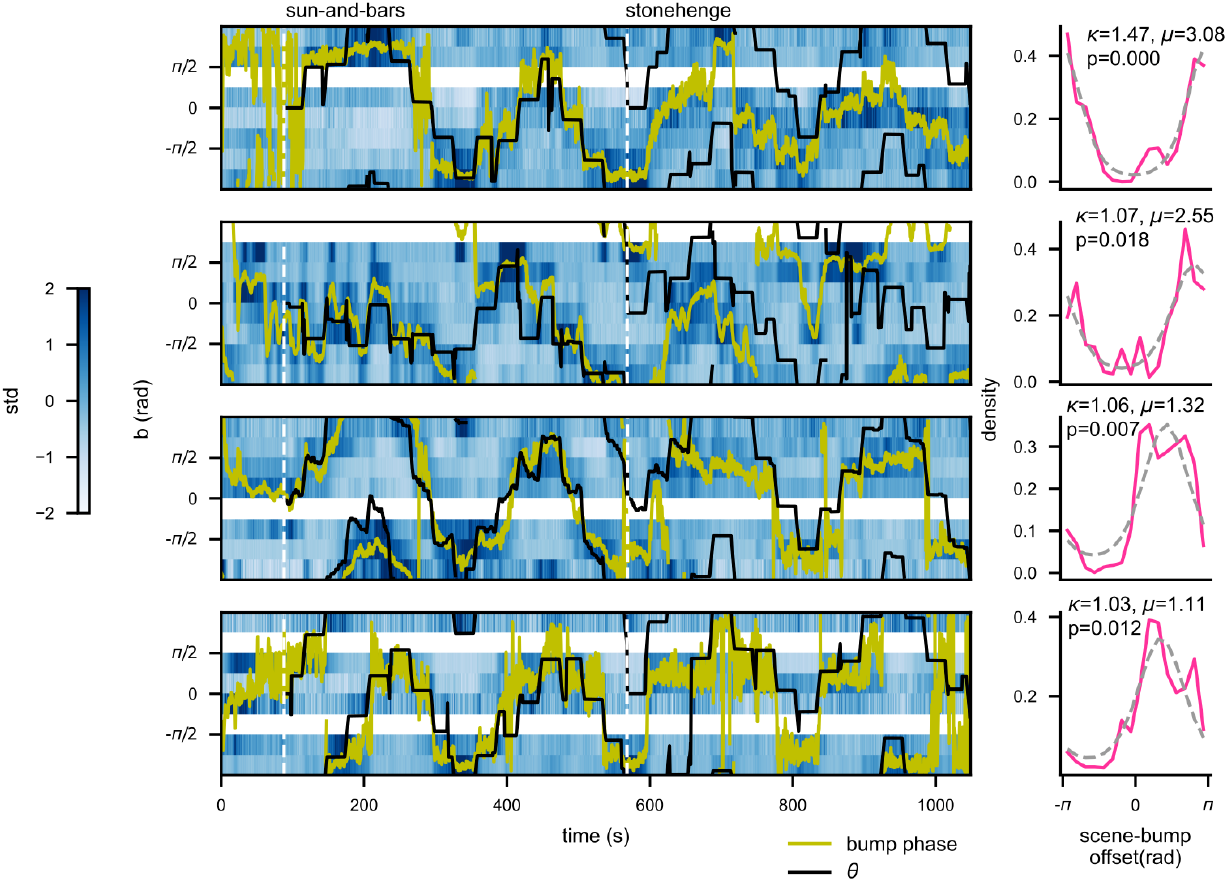
Examples of fish whose HD cells tracked multiple scenes well. Binned activities of the HD cells (left) and the histograms of the offsets between the scene orientation and the bump phase in the Stonehenge epoch (right), as in Figure 2b, c. The parameters of the von Mises distribution fit to the data(dotted grey line) as well as associated bootstrap p-values for κ are noted.

**Figure S5:**
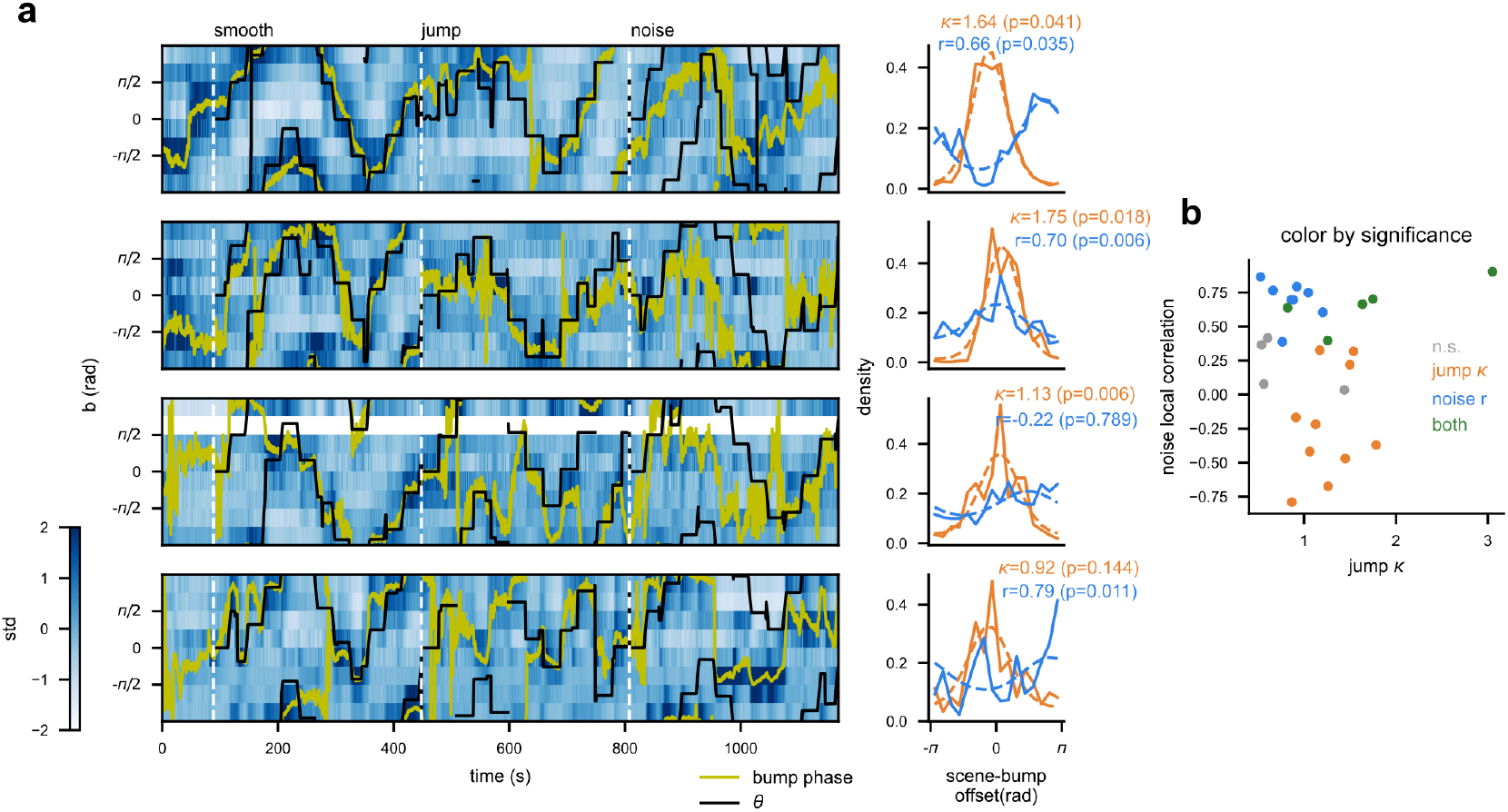
Additional characterizations of landmark- and optic flow-based scene tracking by the HD neurons. **(a)** Binned activities of HD cells with bump phase and θ overlaid(left) and the associated histograms of the scene orientation-to-bump phase offsets (right) from more example recordings, as in Figure 3b, c. The orange and blue solid lines respectively representthe Jumpand Noiseepochs, and dotted lines the von Mises distributions fit on them. The von Mises κ for the Jump epoch and the local correlation for the Noise epoch are indicated with associated p-values from the bootstrap tests. **(b)** The von Mises κ for the Jump epoch plotted against the local correlation in the Noise epoch for each recording, color-coded by the statistical significance of the two metrics. Out of 27 recordings, 5 showed both significant Jump epoch κ and Noise epoch correlation (green), 10 showed only significant κ (orange), 8 showed only significant correlation (blue), and 4 neither (grey).

**Figure S6:**
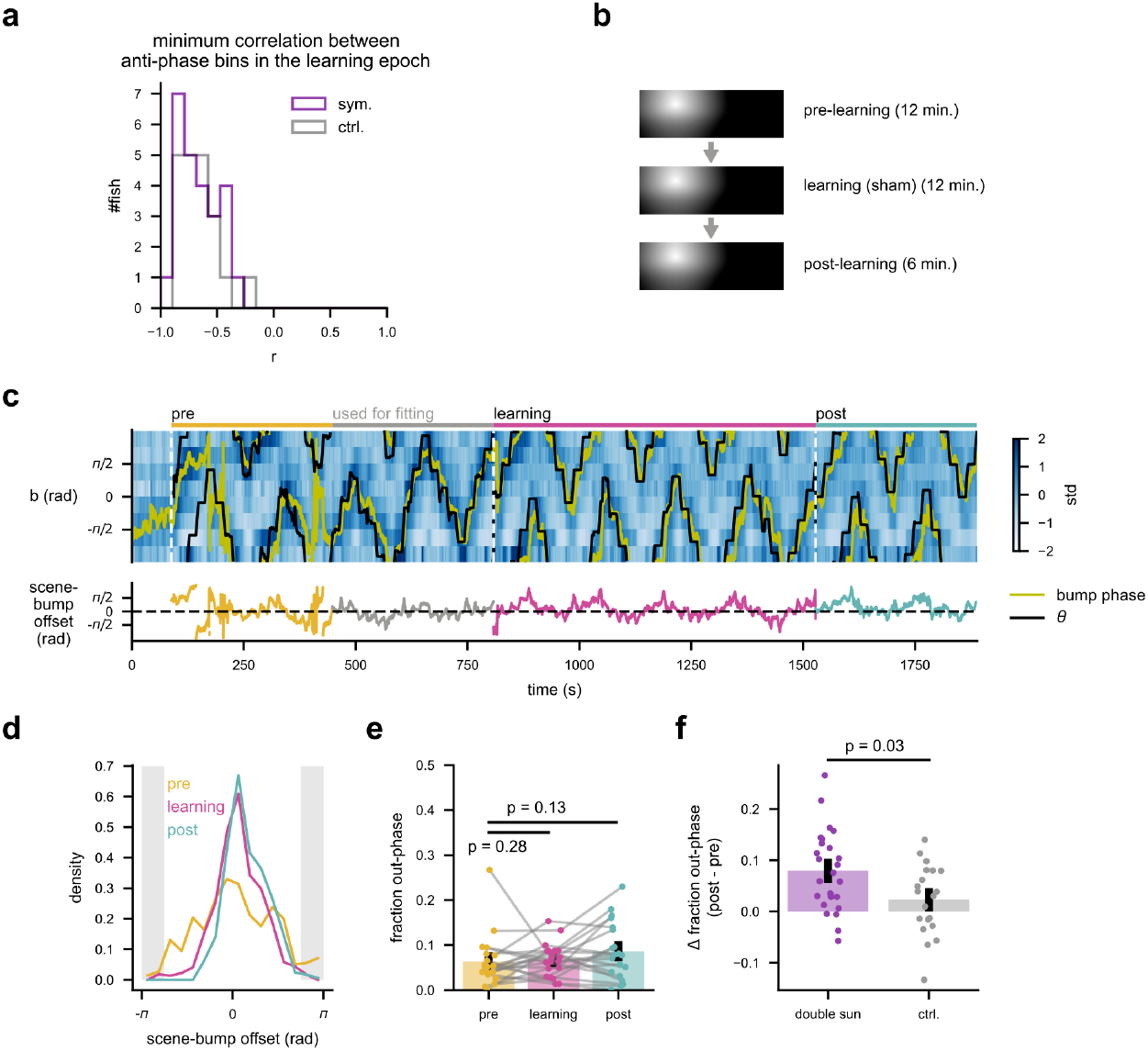
Remapping between the HD neuron bump and the scene orientation does not simply result from the passage of time. **(a)** The distributions of the minimum Pearson correlations between pairs of HD neuron bins with the opposite scene orientation tunings during the learning epochs. Purple and grey lines indicate fish from the double-sun and the control experiment, respectively. The two-fold symmetry in the visual scene would impose strong positive correlation between pairs of visual neurons with azimuthally opposite receptive fields. The fact that these cells remained strongly anti-correlated in the presence of the scene symmetry reveal their strong inhibitory interactions. **(b)** Schematic of the control experiment. **(c)** Data from an example fish in the control experiment, as in Figure 4b. Note how the scene-bump offset remained centered about 0 throughout. **(d)** The distribution of the scene-bump offset for each epoch, for the same fish as shown in (c). **(e)** The fractional time that the bump-scene offset spent in the out-phase range, for each epoch in the control experiment. No significant increase was detected (sign-rank test). N = 20 fish. **(f)** Increase in the fraction of time that the offset spent in the out-phase range from the pretopost-learning epoch was significantly higher in the double-sun experiment, compared to the control experiment (rank sum test, N = 25 fish for the double-sun experiment, 20 fish for the controls).

**Figure S7:**
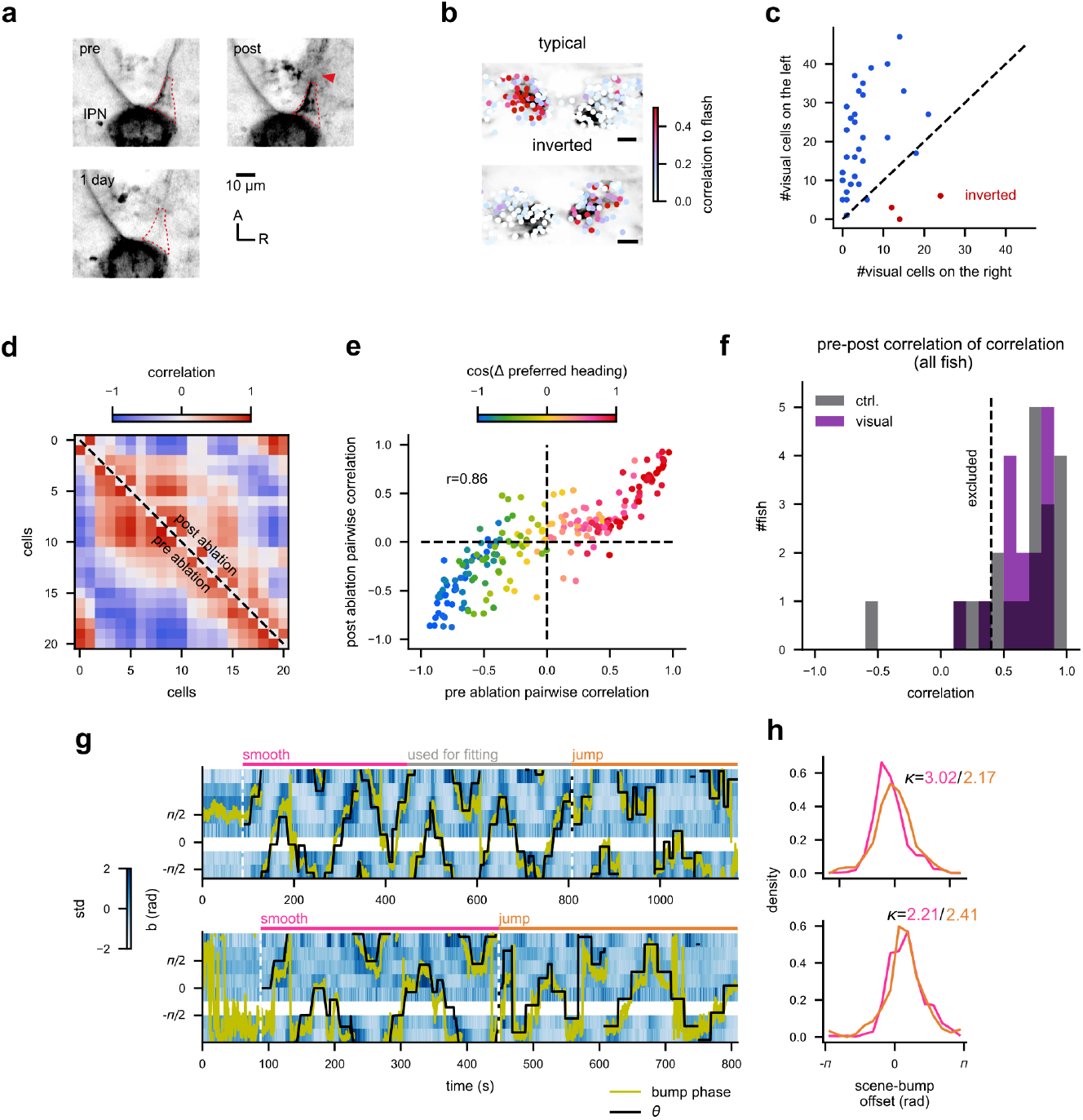
Additional details of the ablation experiment. **(a)** The axons of the habenula neurons entering the IPN, before, right after, and 1 day after the ablation procedure. The ablated site is marked by the red arrow. The neuronal processes between the ablation site and the IPN initially increased the fluorescence, and the disappeared (red dotted line). **(b)** Two examples of fish with the typical and inverted habenula laterality. The ROIs are color-coded by the average pairwise correlations of their responses to the repeated presentations of the flash stimuli. **(c)** The numbers of the habenula ROIs that had averaged flash correlation beyond 0.3 on each side, plotted against each other. The red dots represent fish with the inverted habenula laterality. **(d)** The correlation matrices of the HD neuron activity in the pre-ablation (lower triangle) and post-ablation recordings (upper triangle), in the same example fish as in Figure 5c. The cells are sorted by their preferred scene orientation. **(e)** The pairwise correlation between the all pairs of the HD neurons from the pre- and post-ablation recordings in the same fish, plotted against each other. The cell pairs are colored by the cosine of the difference of their scene orientation tunings. The correlation structure among the HD cells were well maintained in this fish after the ablation. **(f)** The distributions of the correlations of pairwise HD cell correlations before and after the ablations. Fish with correlation lower than 0.4 were excluded from the analysis. **(g, h)** The binned HD cell activities (g) and the scene-bump offset histograms (h) as in Figure 5c, d, but for an example fish with the control side ablation. A: anterior; R: right.

**Figure S8:**
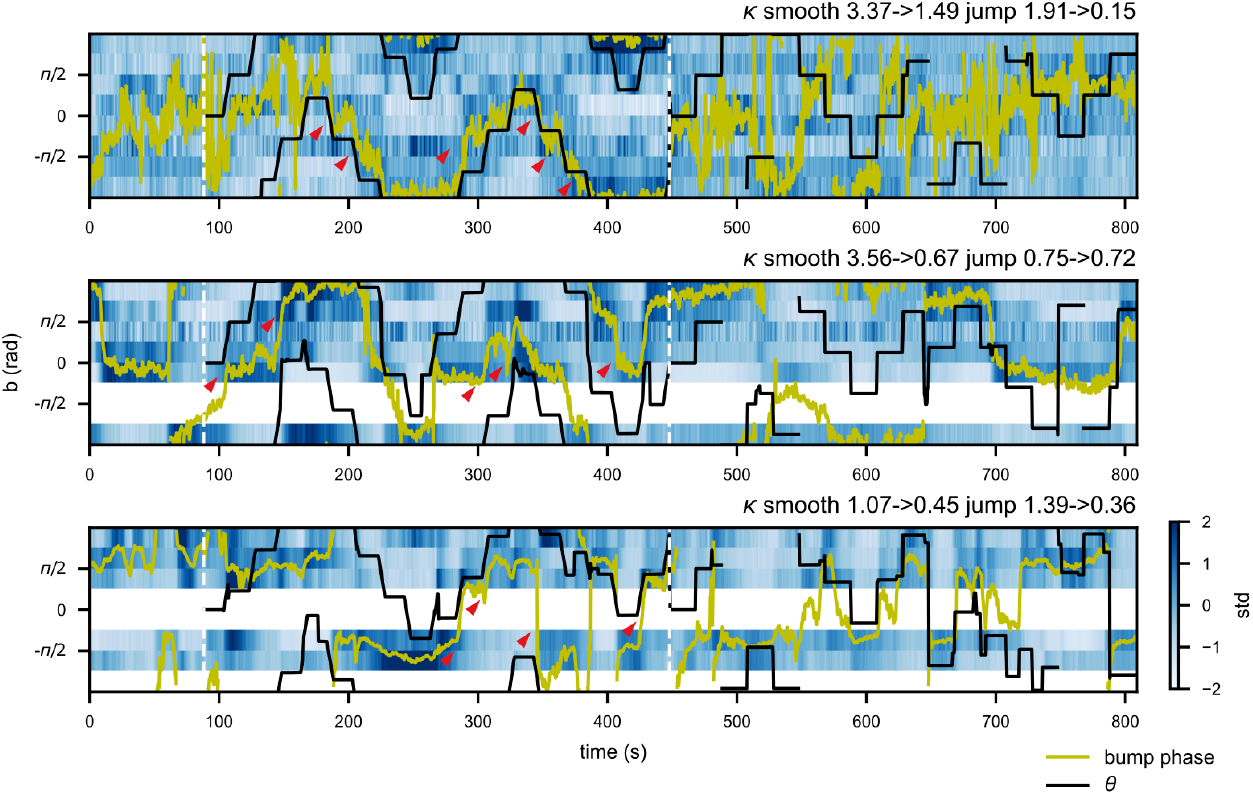
Examples of fish whose bump appeared to move with the scene rotation after the visual side habenula ablation. The post-ablation binned HD cell activities with the scene orientation θ (black) and bump phase (yellow) overlaid, as in Figure 5c. The pre- and post-ablation von Mises κ for each epoch are noted on top. Even though these fish showed overall poor alignment between the scene orientation and the bump phase as indicated by the low postablation κ, the bump phases till seemed to move as the scene rotated in the smooth epoch(the first half of the recording), as indicated by the red arrows. These observations imply that the optic flow information is delivered to the HD cells not from the habenula.

